# MultiPhATE2: Code for Functional Annotation and Comparison of Bacteriophage Genomes

**DOI:** 10.1101/2020.10.05.324566

**Authors:** Carol L. Ecale Zhou, Jeffrey Kimbrel, Robert Edwards, Katlyn McNair, Brian A. Souza, Stephanie Malfatti

## Abstract

To address the need for improved tools for annotation and comparative genomics of bacteriophage genomes, we developed multiPhATE2. As an extension of the multiPhATE code, multiPhATE2 performs gene finding and functional sequence annotation of predicted gene and protein sequences, and additional search algorithms and databases extend the search space of the original functional annotation subsystem. MultiPhATE2 includes comparative genomics codes for gene matching among sets of input bacteriophage genomes, and scales well to large input data sets with the incorporation of multiprocessing in the functional annotation and comparative genomics subsystems. MultiPhATE2 was implemented in Python 3.7 and runs as a command-line code under Linux or MAC-OS. MultiPhATE2 is freely available under an open-source GPL-3 license at https://github.com/carolzhou/multiPhATE2. Instructions for acquiring the databases and third party codes used by multiPhATE2 are found in the README file included with the distribution. Users may report bugs by submitting issues to the project GitHub repository webpage. Contact. Supplementary materials, which demonstrate the outputs of multiPhATE2, are available in a GitHub repository, at https://github.com/carolzhou/multiPhATE2_supplementaryData/.

## Introduction

As the era of reliable antibacterial treatment draws to a close, bacteriophage (phage) therapy is gaining ground in Western countries as an alternative treatment for antibiotic resistant and chronic recalcitrant bacterial infections, with several clinical trials having recently been initiated (Duplessis and Biswas 2020; Pires *et al.* 2020; Altamirano and Barr 2019; Furfaro *et al.* 2018; Gorski *et al.* 2019; Voelker 2019). Next-generation sequencing efforts have generated increasingly large quantities of phage genomic data (Carrol *et al.* 2018; Russell and Hatfull 2017), but still there remains an urgent need for bioinformatics tools for the annotation and evaluation of phage genomic data in support of research efforts toward developing novel phage therapies and therapeutics based on phage products (Gorski *et al.* 2019; Yang *et al.* 2014). A number of advanced bioinformatics tools and computational systems have been developed for evaluation of phage sequence, including detection of prophage sequences within bacterial genomes (Akhter *et al.* 2012; Reis-Cunha *et al.* 2019; and others), evaluation of phage sequence for therapeutic goals (Philpson *et al.* 2018), working with phage metagenomics data (Kleft *et al.* 2020), and microbial genome annotation (Li *et al.* 2016, Seemann 2014). However, most tools used for phage genome annotation are primarily aimed at evaluation of bacterial sequence, and not necessarily suited for use by phage research laboratories with few or no bioinformatics specialists on staff, and limited resources dedicated to constructing annotation and comparative genomics pipelines from individual tools and databases. We developed multiPhATE2 to address this need.

### MultiPhATE2 System Overview

MultiPhATE2 comprises a comprehensive, high-throughput functional annotation and genome comparison system for analysis of newly sequenced phage genomes. MultiPhATE2 performs gene finding followed by computational functional annotation of user-specified phage genomes, then performs gene-by-gene comparisons among the genomes. A system driver script takes a single argument consisting of a configuration file, then invokes up to four computational subsystems: the Gene Calling and PhATE annotation subsystems are run for each genome, and, if two or more genomes are specified by the user, multiPhATE will identify corresponding genes among the genomes using the Compare Gene Profiles (Tkavc *et al.* 2018) and Genomics subsystems (Fig 1).

**Fig. 1.**
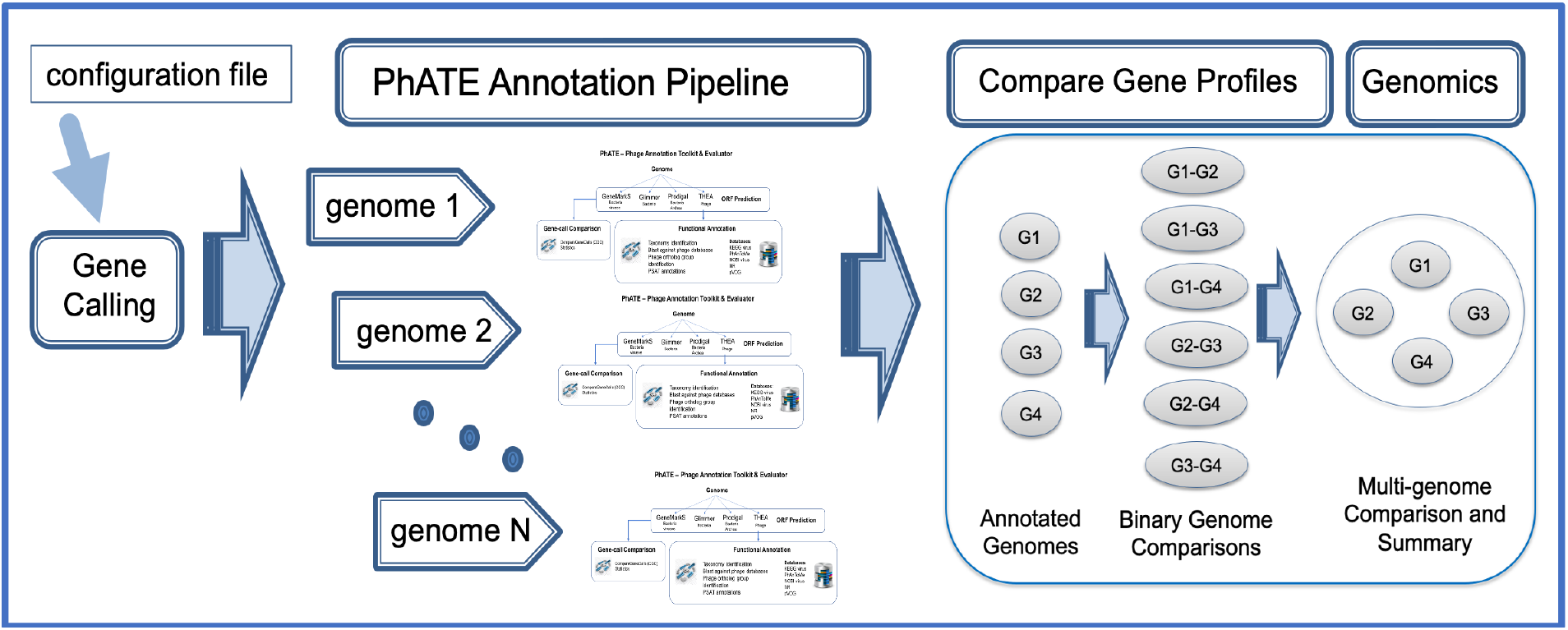
Overview of multiPhATE2 system and workflow. User-specified configurations (configuration file) are input to the multiPhATE2 system, which invokes four subsystems: Gene Calling, the PhATE annotation pipeline, Compare Gene Profiles, which performs binary genome-to-genome comparisons of genes and proteins, and Genomics, which consolidates binary comparisons into gene-gene and protein-protein correspondences among all input genomes.

Based on experiences using multiPhATE2’s predecessor, multiPhATE (Zhou *et al.* 2019), to process input data sets comprising many relatively large phage genomes, we recognized a compelling need for flexible and rapid computations, and we upgraded multiPhATE with this goal: process control features were added for the multiPhATE2 system (Fig. 2). We believe that multiPhATE2 scales well in performing multi-genome annotation and comparison, while offering a flexible toolset for tailoring analyses according to the user’s needs.

**Fig. 2.**
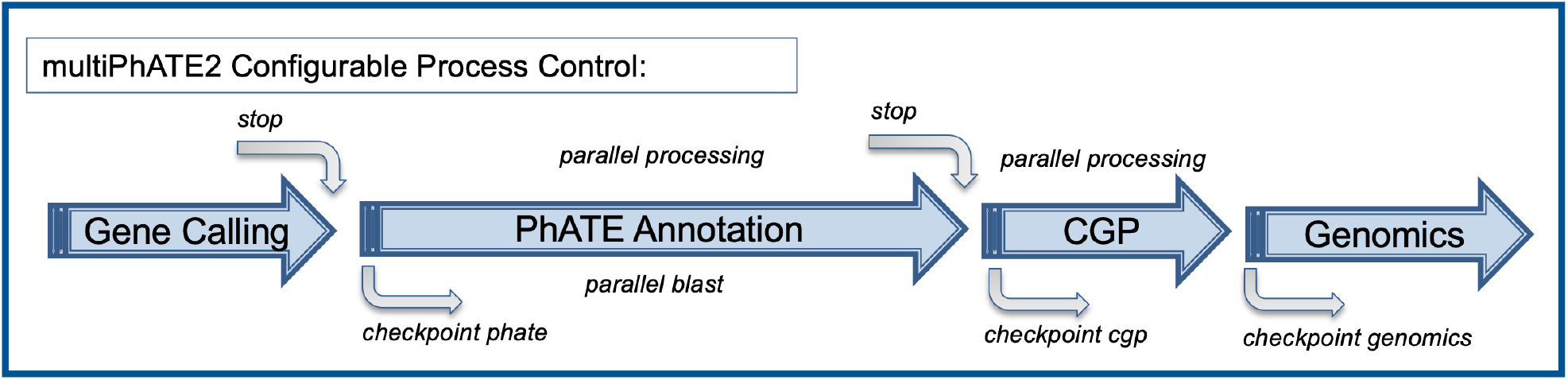
System overview and configurable process control features of multiPhATE2. Large blue arrows: multiPhATE2 subsystems; CGP = Compare Gene Profiles; curved grey arrows: process controls (stop = stopping point; checkpoint = point at which processing may be restarted); “parallel processing” indicates multiprocessing applied to functional annotation of input genomes and binary genome-to-genome comparisons; “parallel blast” indicates multithreading option provided by BLAST+.

## Process Control

Selective and parallel processing are optionally applied at several stages of computation (Fig. 2). These options may be selected in the multiPhATE2 system configuration file. Specifically,

1. Parallel processing is applied in the PhATE and Compare Gene Profiles subsystems. Each genome input to PhATE may be processed as a separate, parallel process, and each binary genome comparison in the Compare Gene Profiles subsystem may be processed likewise in parallel.
2. The user may specify the desired number of threads with which to invoke Blastp (Comacho *et al.* 2009) in the PhATE subsystem.
3. The user may opt to process the multiPhATE2 code through the Gene Finding subsystem or the PhATE subsystem and stop at either point.
4. The user may opt to re-start processing at three points in the multiPhATE2 system. Checkpoints (re-starting points) may be selected at the beginning of the PhATE, Compare Gene Profiles, or Genomics subsystem processing.

### Input

MultiPhATE2 is invoked using a single input parameter consisting of a configuration file, as follows: $ python multiPhate.py multiphate.config. The configuration file allows the user to specify a) genomes to be processed, b) gene finder(s) to be used, c) PhATE annotation search algorithms and databases, d) blast cutoffs, e) locations of databases, f) Compare Gene Profiles matching cutoff, g) PhATE and Compare Gene Profiles multiprocessing, h) stopping points, i) checkpoints, j) and console messaging verbosity. Concise instructions for creating a multiphate configuration file are provided in the project README and in the sample.multiphate.config file provided with the multiPhATE2 distribution. Code execution can be tailored to run specific gene finders and to search for homologous sequences in specific phage- and virus-centric data sets, in addition to more generic protein data sets.

### Gene Calling Subsystem

The Gene Calling subsystem of multiPhATE was updated to include user-provided GFF-formatted custom gene calls, in addition to the already-supported Glimmer (Delcher *et a*l. 2007), GeneMarkS (Besemer *et al.* 2001), Prodigal (Hyatt *et al.* 2010), and PHANOTATE (McNair *et al.* 2019) gene callers, so that it is now also possible to include and compare gene calls from web-only based services, Genbank gene calls, or hand-curated gene-call data sets (see README in the project repository for format specifications). The side-by-side comparison among gene callers (Compare Gene Calls module, Fig. 3) was expanded to include output data sets that either merge the results of multiple gene callers (i.e., a non-redundant superset), or that recognize agreement among callers: a consensus gene-call set, comprising calls that were in agreement among two or more callers, and a common-core gene-call set representing calls that were made by all the gene callers. Any one of the gene-call outputs, custom calls, or multiPhATE-generated gene-call super/subsets may be forwarded on for PhATE functional annotation.

**Fig. 3.**
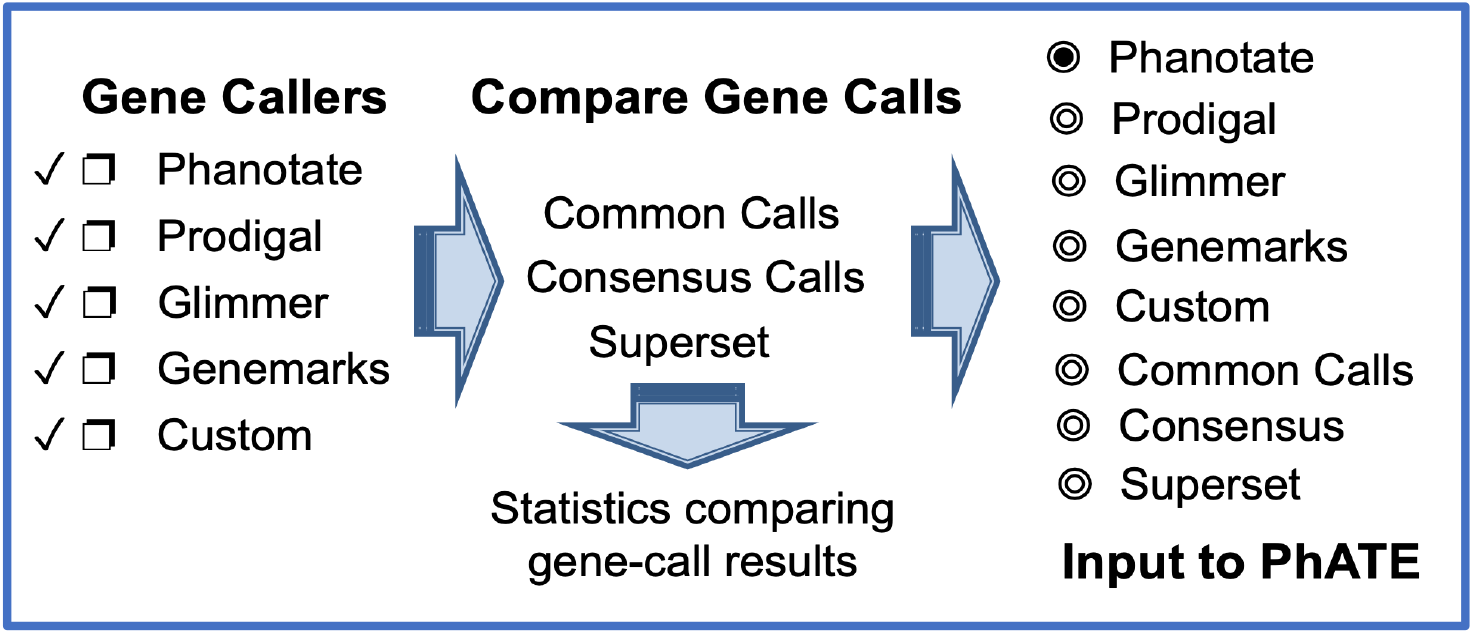
Gene callers and gene-call comparison in the Gene Calling subsystem of multiPhATE2. The user may select any or all of the supported gene callers and/or provide their own gene calls (Custom). The Compare Gene Calls module computes a set of calls that are common among all selected callers (Common Calls), a consensus set comprising gene calls produced by at least two callers (Consensus Calls), and a non-redundant superset of gene calls (Superset). The user may select the results of one gene caller or a super/subset for input to the PhATE subsystem.

### PhATE Annotation Subsystem

PhATE is a fully automated computational pipeline for functional annotation of phage genes within a genome sequence, and was originally written as part of multiPhATE (Zhou *et al.* 2019). Newly incorporated analyses include: blast and hmm searches for VOG gene and protein (Laffy *et al.* 2016), CAZy (Lombard *et al.* 2014), custom genome, gene, and protein databases; HMM profile searches for pVOG-hmm (Grazziotin *et al.* 2017) and VOG-protein-hmm (Laffy *et al.* 2016) databases; and new searches with phmmer and hmmscan (Johnson *et al.* 2010). Preprocessing of the pVOG, VOG, and CAZy data sets are performed in order to speed the matching of hits with associated annotations. A script is provided to facilitate downloading, preprocessing, and updating of databases that are supported in multiPhATE2. A full accounting of the search algorithms and databases supported by multiPhATE2 is depicted in Fig. 4

**Fig. 4.**
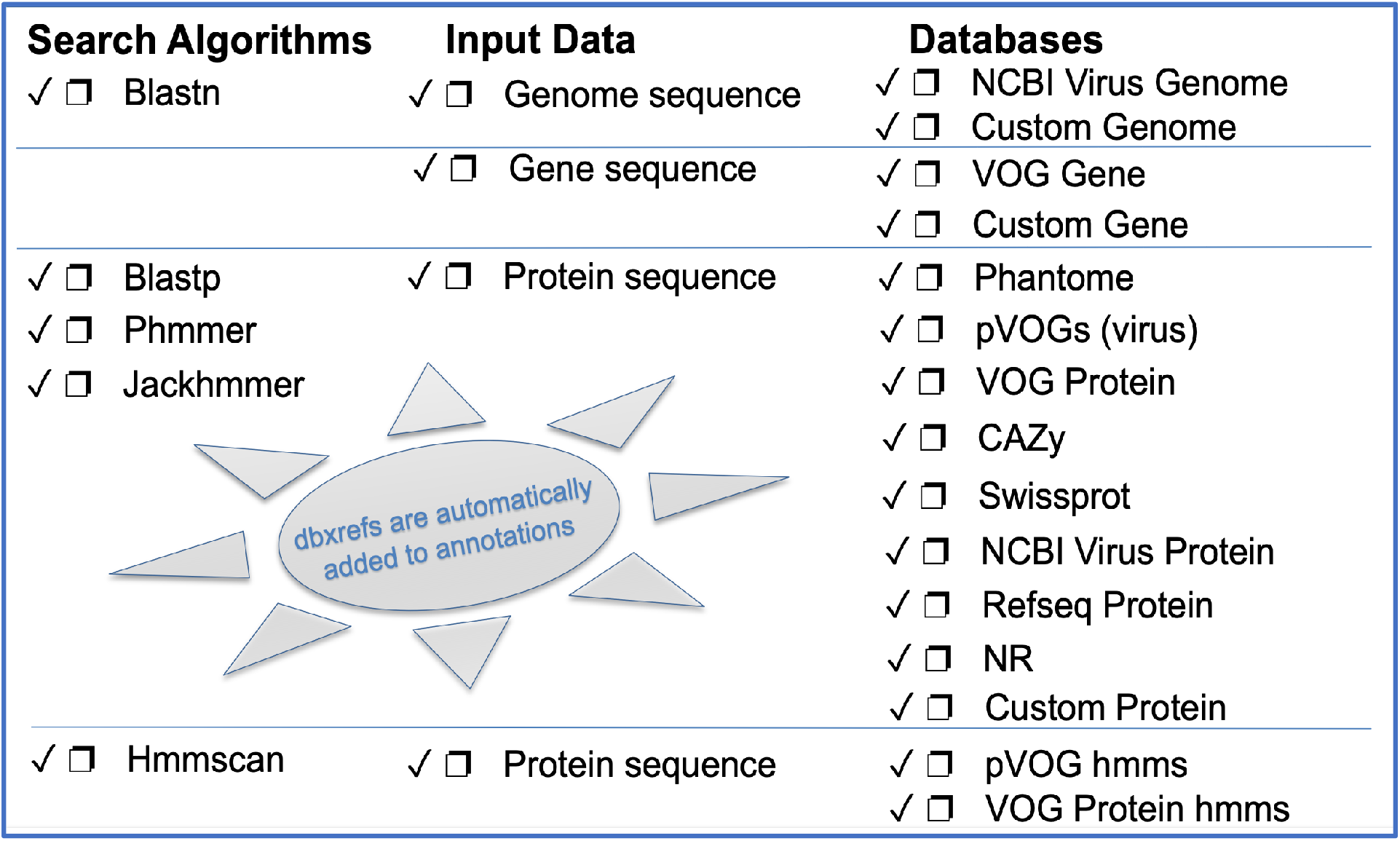
Functional annotation options supported within the PhATE subsystem of multiPhATE2. The user may select any or all of the algorithms to search any or all of the databases over genome, gene, and/or protein sequences.

### Comparative Genomics Subsystems

MultiPhATE2 accomplishes comparisons among input genomes in a two-step process whereby each genome is compared to each other, and then gene-gene and protein-protein correspondences are identified from among all of the input genomes.

The Compare Gene Profiles subsystem performs NxN reciprocal blast of the genes from each genome against the genes from every other genome provided by the user (Fig. 1). The code then identifies for each gene its mutual and non-mutual (singular) best hits against corresponding genes from each of the other genomes, or reports if no corresponding gene is found. For each binary genome-to-genome comparison, hits are ordered with respect to the query genome. The Genomics subsystem inputs the binary blast result files from Compare Gene Profiles and computes genes and proteins that correspond across all the input genomes with respect to the reference genome (first genome listed). Ultimately, homology groups are generated, comprising each reference gene and its corresponding genes, plus its paralog’s corresponding genes. This analysis is also performed for protein sequences. PhATE annotations are carried through the Compare Gene Profiles and Genomics computations so that the user can readily identify gene/protein function among the identified homolog groups.

### MultiPhATE System Output

Directories and files that are produced by multiPhATE2 are detailed in the README. In brief, an output subdirectory is created for each input genome to hold results of the Gene Finding and PhATE subsystems, and subdirectories are created to hold results of the Compare Gene Profiles and Genomics subsystems. A sample multiPhATE2 system main output directory with full contents can be found in the multiPhATE2 supplementary data repository. In summary, the following subdirectories are created:

1. Genome result directories, one for each genome processed through the Gene Calling and PhATE subsystems, including a) results of gene calling, b) BLAST, HMM, and PROFILE directories containing intermediate and final results of searches, c) fasta groupings comprising query sequences with their pVOG or VOG homologs, d) final results of side-by-side gene calling comparison (file: CGC_results.txt) and final results of functional annotation (files: phate_sequenceAnnotation_main.out/.gff).
2. CGP_RESULTS directory: Output from the Compare Gene Profiles subsystem,
3. GENOMICS_RESULTS directory: Output from the Genomics subsystem.
4. JSON directory: Contains one automatically generated JSON configuration file for each instance of PhATE annotation (*i.e.,* per genome). The JSON files convey user-specified input parameters to the PhATE subsystem.

## Summary

MultiPhATE2 represents a significant advance in bacteriophage genome annotation in that it streamlines gene calling, functional sequence annotation, and comparative genomics for sets of newly sequenced draft or finished genomes. MultiPhATE is straight forward to install, with full instructions in the README file in the project’s github repository. Running multiPhATE2 as a command-line program taking a single argument (*i.e.,* the multiphate.config file) facilitates launching jobs comprising annotation and comparison of potentially large sets of genomes. Furthermore, built-in flexibility (Fig. 2) allows the user not only to install components in a step-wise manner, but also to run and re-run each subsystem with different parameters, so that the user may determine, for example, a) which gene caller or callers are preferred and which one may be best to carry through to functional annotation; b) which search algorithms and databases (including custom) are most appropriate for the genomes under study; c) which sequence identity cutoffs produce the desired stringency in terms of homolog identification or gene-gene correspondence. MultiPhATE2 offers a wide range of gene calling and sequence search algorithms and databases, including the options of providing user-defined custom gene calls and blast databases (Figs. 3, 4). Parallel processing within the PhATE and CGP subsystems (Fig. 2) enables computations for large input data sets, which might not otherwise be feasible when processing in serial. Furthermore, hits to entries in each of the pVOG and VOG databases are combined with the query protein sequence to produce fasta groupings to facilitate potential follow-on analyses (beyond the scope of multiPhATE2), such as multiple sequence alignment and common motif identification. We know of no other phage-tailored genome annotation system that provides breadth and flexibility that are comparable to multiPhATE2.

## Acknowledgments

This work was performed under the auspices of the US Department of Energy by Lawrence Livermore National Laboratory under contract DE-AC52-07NA27344.

## Funding

This work was supported by the U. S. Department of Defense Defense Threat Reduction Agency grant number DTRA13081-32220 and the U. S. Department of Energy Office of Biological and Environmental Research award numbers SCW1632 and SCW1656.

## Conflicts of interest

None declared.

## Notes

https://github.com/carolzhou/multiPhATE2/

https://github.com/carolzhou/multiPhATE2_supplementaryData

### Competing Interest Statement

The authors have declared no competing interest.

